# Deletion of Setd7 protects against cardiac hypertrophy via inhibiting lipid oxidation

**DOI:** 10.1101/2024.07.29.605718

**Authors:** Haibi Su, Jinghuan Wang, Yuyu Zhang, Jie Xu, Jiayao Liu, Yuhui Li, Chenxi Xiao, Caiyun Wang, Jun Chang, Xinhua Liu

## Abstract

Setd7, a catalytic enzyme responsible for histone H3K4 methylation, is implicated in various cardiac diseases. However, the role of Setd7 in pathological cardiac hypertrophy remains unclear. In this study, we observed that Setd7 is significantly elevated in pathological hypertrophy stimuli cardiomyocytes and mouse failing hearts. Subsequently, we found that mice lacking Setd7 remarkably preserved cardiac function after transverse aortic constriction, as demonstrated by improving myocardial hypertrophy and fibrosis, whereas Setd7 overexpression in cardiomyocytes deteriorated hypertrophy phenotype. Further in vitro analyses revealed that Setd7 mediated-E2F1 activation induces E3 ubiquitin protein ligases WWP2 expression to catalyze the lipid-peroxide-reducing enzyme GPx4 ubiquitination degradation, ultimately causing widespread lipid peroxidation and boosting pathological cardiac hypertrophy. Remarkably, loss of activity of GPx4 blunted the Setd7 knockdown exerts antihypertrophic effect in pathological cardiomyocytes hypertrophy, further confirming an important role of lipid peroxidation in Setd7-mediated failing hearts. In summary, the role of Setd7 in pressure overload-induced cardiac hypertrophy is regulated by the Setd7-E2F1-WWP2-GPx4 signaling pathway, suggesting that targeting Setd7 is a promising therapeutic strategy to attenuate pathological cardiac hypertrophy and heart failure.

## Introduction

Pathological cardiac hypertrophy occurs in response to volume or pressure overload, ultimately leading to contractile dysfunction and increased risk for heart failure (HF) (Groenewegen *et al*, 2020) (Dorn *et al*, 2003; Schiattarella & Hill, 2015). Sustained cardiac hypertrophy is usually accompanied by interstitial and perivascular fibrosis and cardiomyocyte death, with increased levels of collagen and myofibroblast activation. Accumulating evidence suggests that mechanical stretching, oxidative stress, and inflammation mediated molecular mechanisms promote maladaptive cardiac remodeling and dysfunction. Thus, finding the specific regulatory targets to delay or reverse cardiac hypertrophy might provide new therapeutic strategies for preventing HF.

Lipid peroxidation (LPO) is a process of lipid electrons in the cell membrane removed by free radical species such as oxyl radicals, peroxyl radicals, and hydroxyl radicals(Yin *et al*, 2011). This process subsequently generates overproduction of reactive oxygen species (ROS), resulting in the oxidation of membrane lipids polyunsaturated fatty acids fatty acids (PUFAs) to form lipid hydroperoxides (LOOH) (Gaschler & Stockwell, 2017). Low rates of LPO are under cellular control, but when some threshold of LPO is reached, and get out of control, thus driving the disruption of membrane integrity and function. Lipid peroxidation can be combated by glutathione peroxidase 4 (GPx4), a main endogenous antioxidant peroxidase that directly reduces phospholipid hydroperoxide(Chen *et al*, 2024a). GPx4 activity is essential to maintain lipid homeostasis in the cell, inhibition of GPx4 function leads to lipid peroxidation, the ROS accumulation, and can trigger ferroptosis, an iron-dependent cell death(Dixon *et al*, 2012; Stockwell *et al*, 2017). Lipid peroxidation, a prevalent feature of a number of disease states, has become an important mechanism in the development of cardiovascular diseases, including pathological cardiac hypertrophy (Zhang *et al*, 2023).

Numerous reports strongly argue that epigenetic modifiers play crucial roles in the development of various diseases and pathological processes. The SET domain-containing lysine methyltransferase Setd7 methylates histone 3 lysine 4 (H3K4) in the enhancer or promoter regions associated with transcriptional activation. Recently, a number of non-histone targets such as STAT3, β-catenin, Hif-1α, etc. were shown to be methylated by Setd7(Liu *et al*, 2015; Lv *et al*, 2023; Oudhoff *et al*, 2016). Therefore, Setd7 may function mainly through the transcriptional activation or methylation of different non-histones, which regulate various cellular responses, including altered gene expression, protein stability, and loss of DNA methylation. While Setd7 has been well studied in different cancers and fibrosis, additional studies also demonstrate Setd7 methylated the Hippo effector YAP drives myocardial ischemic injury(Ambrosini *et al*, 2023). Given the important role of Setd7 in heart, whether Setd7 modulates the pathogenesis of cardiac hypertrophy remains unknown.

To determine the role of Setd7 and Setd7-associated lipid peroxidation in cardiac hypertrophy, we generated mice lacking Setd7. We found that these mice protect cardiac dysfunction and myocardial remodeling from pressure overload. We demonstrated that Setd7 causes GPx4 degradation by directly enhancing E3 ubiquitin ligase WWP2 expression via enriching the transcriptionally activation marks H3K4 on the E2F1 promoter, ultimately causing widespread lipid peroxidation and boosting pathological cardiac hypertrophy. Our results elucidate the critical role played by Setd7 in the pressure overload/hypoxia-induced cardiomyopathy and may provide therapeutic targets for the development of improved treatment strategies.

## Materials and methods

### Cardiac primary cell isolation, culture and treatment

Primary myocytes were isolated from 1-3-day-old Sprague-Dawley (SD) rats or *Setd7*^+/+^ and *Setd7*^−/−^ mice. The hearts were quickly removed and washed with PBS and then minced. The minced hearts were digested with a mixed solution of 0.25% trypsin and 1% collagenase II at 37°C. After collecting the digested cell suspension and centrifuging, the cells were resuspended in DMEM containing 10% FBS and 1% penicillin-streptomycin. The cells were pre-plated for 1.5-2 h to remove the attached fibroblasts, and then the cardiomyocytes were cultured at 37°C with 5% CO_2_.

Primary rat cardiomyocytes were plated in plates, and when the cardiomyocytes reached approximately 70% confluence, the cells were transferred to serum-free medium and cultured for 6-12 h. To induce hypertrophy, the cells were treated with 100 μM phenylephrine (PE) for 48 h. In addition, a hypoxic incubator was used with a gas concentration of 1% O_2_, 5% CO_2_, and 94% N_2_ to induce hypoxia for 6 h. The hypertrophic response was evaluated by measuring cell surface area and the expression of hypertrophic markers.

### Hypobaric hypoxia model

Eight-week-old male C57BL/6 mice were subjected to hypobaric hypoxic environment by an environment simulative cabin simulating an altitude of 6000 m (9% O_2_) for 8 weeks. Hypertrophic markers were measured to evaluate ventricular hypertrophy.

### Transverse aortic constriction

Transverse aortic constriction (TAC) was performed on 8-10-week-old male C57BL/6 mice to induce cardiac hypertrophy. Briefly, mice were anesthetized with 1.5% isoflurane and performed endotracheal intubation. A left sternotomy was performed to expose the aorta arch. A 7-0 silk suture was passed through the aortic arch between the innominate artery and the left common carotid artery, and a 27-gauge needle was placed parallel to the aortic arch. The suture was then tied around both the needle and the aorta to create the ligation, and the needle was subsequently removed to create the constriction. Sham-operated mice underwent the same procedure but without aortic constriction.

### Statistical analysis

Data were expressed as mean ± SD. Differences of means were analyzed by using one-way ANOVA with the Tukey-Kramer post hoc test for multiple groups, and when comparing between two groups using unpaired Students *t*-test. All analyses were made using the *GraphPad Prism 7.0* statistical software package and *p* value of < 0.05 was accepted as statistically significant.

## Results

### Hypertrophic stimulation results in an increase in the expression of Setd7 in vivo and vitro

Using differently pathological cardiac hypertrophy model, we first explored Setd7 expression during cardiac hypertrophy. Firstly, neonatal rat cardiomyocytes (NRCMs) exposed to 1% O_2_ for 6 h (hypoxia) or treated with potent hypertrophic stimulus phenylephrine (PE) for 48 h showed significantly increased the Setd7 mRNA and protein levels (**Figure 1A-D**). Consistently, hypobaric hypoxia (HH) mice (8 weeks) showed significant right ventricular hypertrophy **(Figure 1E**), and Setd7 mRNA and protein expression were markedly increased relative to normoxia (NN) mice (**Figure 1F-G**). In addition, TAC-challenged mice showed significant left ventricular hypertrophy (**Figure 1H**) and higher Setd7 mRNA and protein levels (**Figure 1I-J**). The protein levels of ANP (atrial natriuretic peptide) and BNP (brain natriuretic peptide), the biomarkers for cardiac hypertrophy, were remarkably increased in hypoxia or PE stimulated myocytes, as well as mice hearts under hypobaric hypoxia and TAC compared with control (**Figure 1A-I**). Similarly, PE or hypoxia induced a near-doubling in size (indicated as the cell-surface area in myocytes) **(Figure 1K).** Collectively, these data consistently demonstrate that Setd7 expression can be regulated by hypertrophic challenge.

**Figure 1.**
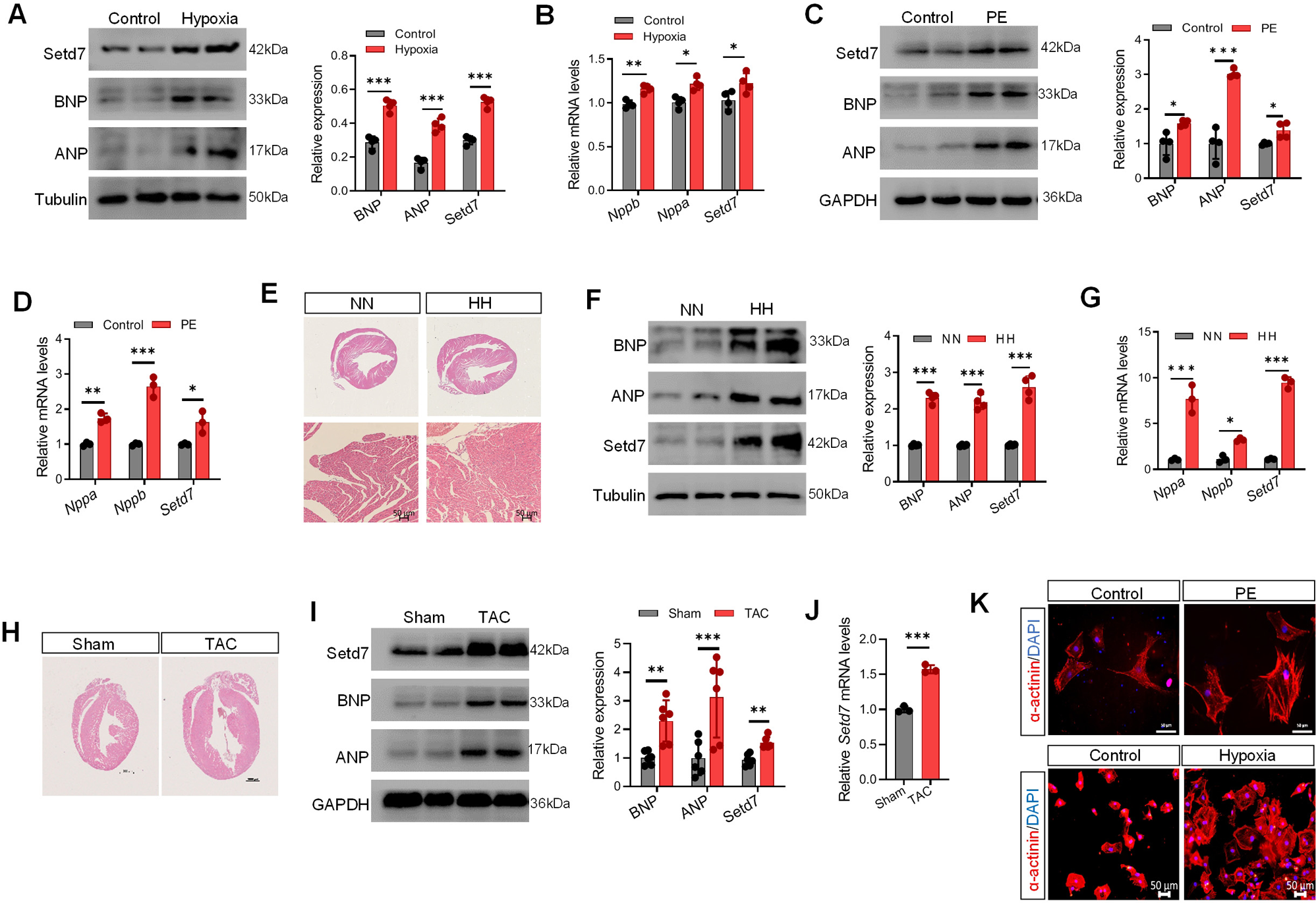
Hypertrophic stimulation results in an increase in the expression of Setd7 in vivo and vitro. (**A**) Neonatal rat cardiomyocytes (NRCMs) exposed to 1% O_2_ for 6 h (hypoxia), protein expressions of BNP, ANP and Setd7 was examined by Western blot and quantitative analysis. (**B**) mRNA levels of *Nppb*, *Nppa*, and *Setd7* were measured by RT-qPCR in normoxia and hypoxia cardiomyocytes. (**C**) The protein expressions of Setd7 and myocardial hypertrophic markers (ANP, BNP) in 100 µM phenylephrine (PE)-induced NRCMs for 48 h were detected by Western blot. (**D**) The mRNA levels of *Setd7*, *Nppa* and *Nppb* in PE-treated NRCMs were detected by RT-qPCR. (**E**) Heart tissue sections stained with hematoxylin-eosin (H&E) from mice exposed to normoxia (NN) and hypobaric hypoxia (HH). (**F**) Protein levels of BNP, ANP, and Setd7 were detected by Western blot in normoxia and hypobaric hypoxia mice. (**G**) mRNA levels of *Nppa*, *Nppb*, and *Setd7* were measured by RT-qPCR in normoxia and hypobaric hypoxia mice. (**H**) Representative images of hearts from mice subjected to sham or transverse aortic constriction (TAC) surgery by H&E staining. (**I**) The protein expressions of Setd7 and hypertrophic markers (ANP, BNP) in heart tissues from mice subjected to sham or TAC surgery were detected by Western blot. (**J**) The mRNA level of *Setd7* in mice subjected to sham or TAC surgery. (**K**) Representative images of α-actinin in PE or hypoxia induced hypertrophic NRCMs. Scale bar: 50 μm. n = 3-6. * represents *p* < 0.05, ** represents *p* < 0.01, *** represents *p* < 0.001 (Student’s t-test).

### Lacking Setd7 mitigates TAC-induced pathological cardiac hypertrophy and fibrosis

To dissect the functional role of Setd7 in pathological cardiac hypertrophy in vivo, *Setd7* heterozygous (*Setd7*^+/−^) mice were generated, Setd7 knockdown in heart tissues from *Setd7*^+/−^ mice was verified by Western blotting and RT-qPCR (**Figure S1**). We subjected WT and *Setd7*^+/−^ mice to 4 weeks of pressure overload and determined cardiac function by echocardiography. Notably, the *Setd7*^+/−^ mice did not exhibit marked abnormalities in heart structure or function under basal conditions. However, TAC-operated *Setd7*^+/−^ mice exhibited remarkably improved cardiac dilation and dysfunction compared with the WT mice, as evidenced by echocardiograph parameters, e.g., left ventricular ejection fraction (LVEF), left ventricular fraction shortening (LVFS), left ventricular end-diastolic and end-systolic internal diameter (LVID), and LV volume (**Figure 2A**). Furthermore, the *Setd7*^+/−^ mice showed significantly decreased ratios of heart weight to body weight (HW/BW), HW to tibia length (HW/TL) compared with WT mice 4 weeks after TAC surgery (**Figure 2B**). Moreover, the grossly apparent differences, along with the haematoxylin and eosin (H&E) and wheat germ agglutinin (WGA) staining, demonstrated that the hearts of *Setd7*^+/−^ mice were not only smaller in size but also had a reduced surface area compared to WT mice (**Figure 2C**). Meanwhile, masson’s trichrome staining and subsequent quantification of the fibrotic area (blue staining) revealed significantly more fibrosis in the hearts of WT mice compared to *Setd7*^+/−^ mice at 4 weeks after TAC (**Figure 2C**). Consistently, the protein and transcript levels of several hypertrophic marker genes (*Nppa*, *Nppb* and *Myh7*) and fibrotic markers (α-SMA, MMP9, *Tgfβ1*, *Mmp2* and *Col1a1*) were decreased with different degrees in the *Setd7*^+/−^ mice compared with the WT mice 4 weeks after TAC surgery (**Figure 2D and E)**. These data demonstrate that Setd7 is involved in regulating the pathologic hypertrophic response of the myocardium to pressure overload stimulation. Moreover, Setd7 ablation plays a protective role against cardiac hypertrophy.

**Figure 2.**
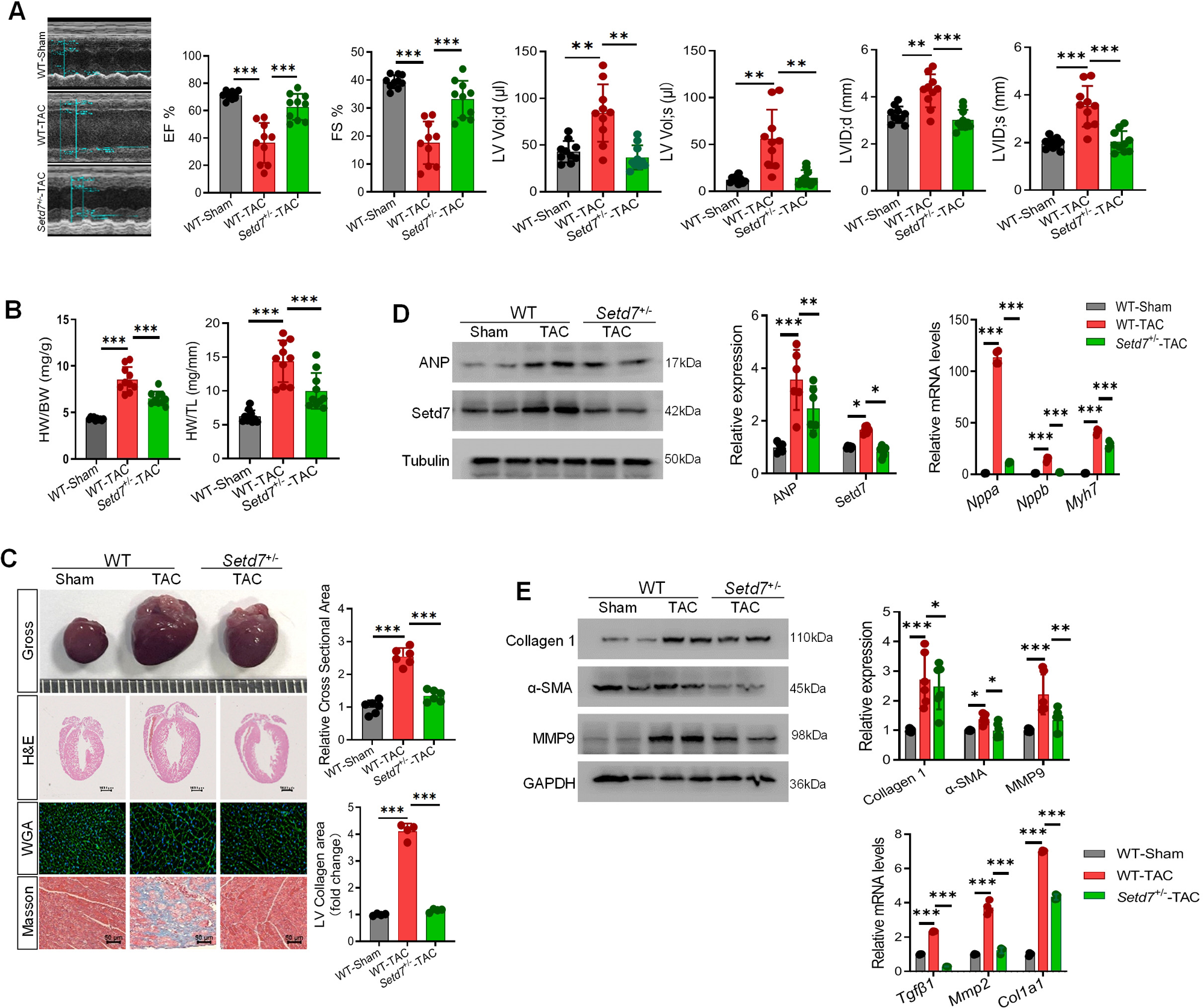
Genetic deletion of Setd7 protects against TAC-induced cardiac hypertrophy. Setd7 heterozygous mice (*Setd7*^+/−^ mice) were subjected to Sham or transverse aortic constriction (TAC) surgery and harvested after 4 weeks. (**A**) Representative echocardiography echo images and EF%, FS%, LVID;d, LVID;s, LV Vol;d and LV Vol;s of wild-type (WT) and *Setd7*^+/−^ mice as determined (n = 10). (**B**) Heart weight normalized by body weight or tibia length of WT and *Setd7*^+/−^ mice after TAC surgery. n = 10. (**C**) Representative images of heart sections stained with H&E, WGA or Masson. Quantitative analysis is presented on the right. n = 4 or 6. (**D**) The protein expressions of Setd7 and ANP, and mRNA levels of *Nppa*, *Nppb*, and *Setd7* were detected by Western blot and RT-qPCR. n = 4 or 6. (**E**) Protein expression (Collagen I, α-SMA, MMP9) and mRNA levels (*Tgfβ1*, *Mmp2* and *Col1a1*) of cardiac fibrosis-related gene in the hearts. n = 4 or 6. * represents *p* < 0.05, ** represents *p* < 0.01, *** represents *p* < 0.001 (One-way ANOVA).

### Blocking Setd7 suppresses cardiomyocyte hypertrophy

To confirm the link between Setd7 and pathological cardiac hypertrophy, we next examined whether it could also regulate the hypertrophic response in cultured neonatal cardiac myocytes. To this end, we lowered endogenous Setd7 expression with a specific siRNA. *Setd7*-specific siRNA transfection significantly reduced the expression in pathological hypertrophic marker gene (ANP, BNP, Myh7) (**Figure 3A and B**) and cell size (**Figure 3C**) upon hypoxia exposure. Meanwhile, we found Setd7 knockdown decreased myocytes inflammatory responses induced by hypoxia (**Figure S2A**). To independently verify the above observation, we isolated neonatal cardiomyocytes from *Setd7*^+/+^ and *Setd7*^+/−^ mice, then cardiomyocytes were exposed to hypoxia. Similarly, immunofluorescent staining for α-actinin demonstrated that Setd7 knockdown reduced hypoxia-induced hypertrophy of cardiomyocytes (**Figure 3D**), accompanied with reduced protein expression levels of ANP and BNP (**Figure 3E**).

**Figure. 3.**
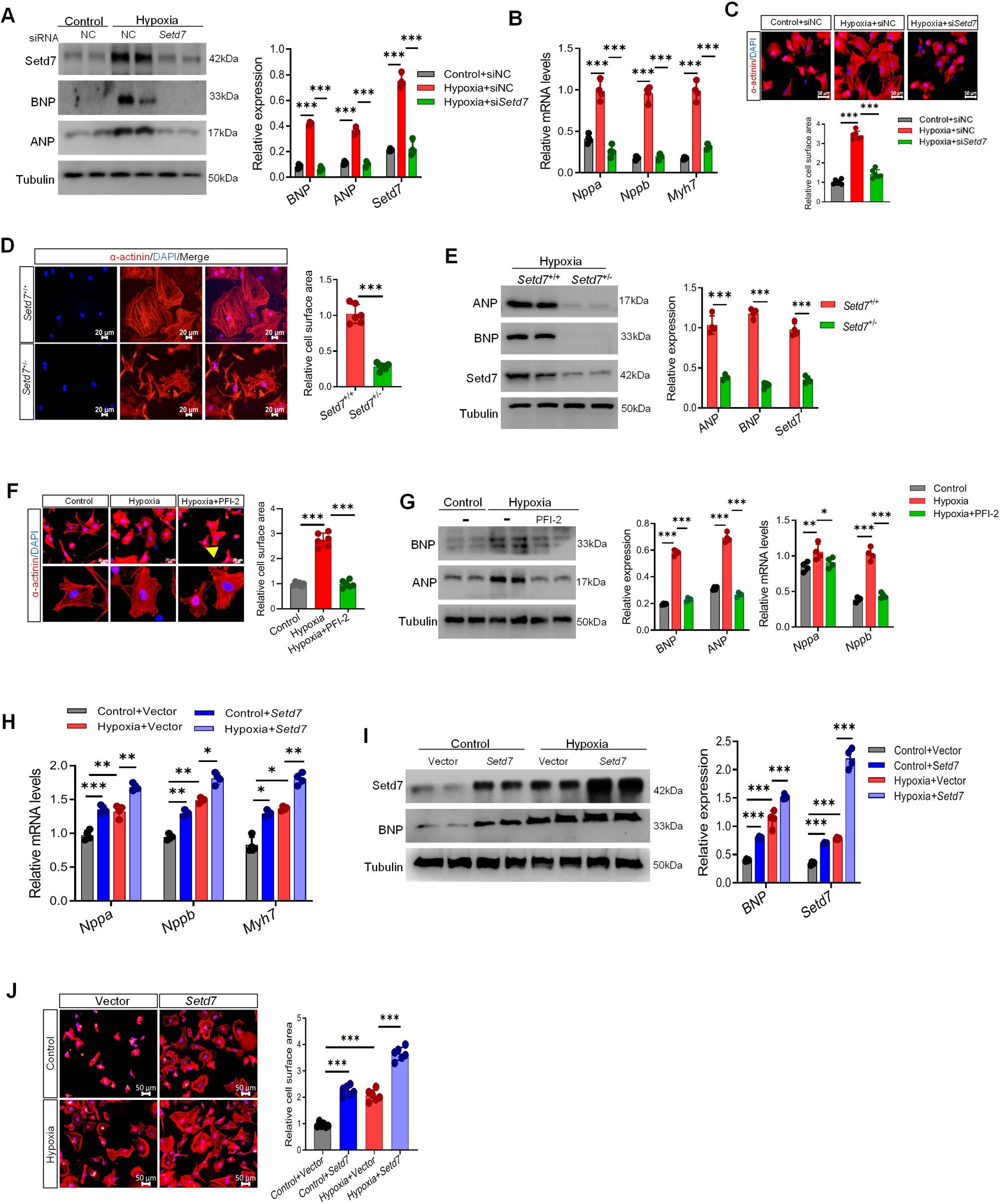
Effect of Setd7 on hypoxia induced hypertrophic NRCMs through function of gain and loss. (**A-C**) NRCMs were transfected with either control siRNA (NC) or Setd7 siRNA and then treated with 1% O_2_ for 6 h to simulate hypoxia. Protein levels of BNP, ANP, and Setd7 were detected by Western blot. (**A**); mRNA levels of *Nppa*, *Nppb* and *Setd7* were detected by RT-qPCR. (**B**); Immunofluorescence staining of α-actinin and cell size was analyzed. Scale bars: 20 μm. (**C**). (**D**) Representative immunofluorescence staining of α-actinin and cell size was analyzed in neonatal *Setd7*^+/+^ and *Setd7*^+/−^ mice NMCMs after hypoxia. Scale bars: 20 μm (**E**) Immunoblot analysis of cardiac hypertrophy associated proteins (ANP and BNP) and Setd7 in neonatal WT and *Setd7*^+/−^ mice NMCMs after hypoxia. (**F**) NRCMs were incubated with PFI-2 and then treated with 1% O_2_ for 6 h to simulate hypoxia. Representative immunofluorescence staining of α-actinin and cell size was analyzed. Scale bars: 20 μm (**G**) Immunoblot analysis of BNP and ANP. (**H**) RT-qPCR to quantify mRNA level of *Nppa*, *Nppb* and *Myh*7 in *Setd7* overexpressing NRCMs with or without hypoxia. (**I**) Immunoblot analysis of Setd7 and BNP in *Setd7* overexpressing NRCMs under normoxia or hypoxia condition. (**J**) Immunofluorescence staining of α-actinin in *Setd7* overexpressing NRCMs. Scale bars: 50 μm. n=4 or 6. * represents *p* value < 0.05, ** represents *p* value < 0.01, *** represents *p* value < 0.001 (Student’s t-test or One-way ANOVA).

Next, we analyzed if inhibition of Setd7 can protect hypoxia-induced cardiomyocyte hypertrophy. PFI-2, a pharmacological inhibitor of Setd7 was used, and found PFI-2 blocked hypoxia-dependent hypertrophy of NRCMs (**Figure 3F**) and reversed the prototypical gene program associated with pathological cardiac hypertrophy (**Figure 3G**), as well as inflammatory mediators (**Figure S2B**). Together, these data support the view that Setd7 contributes to hypoxia-induced cardiomyocyte hypertrophy, and blocking Setd7 suppresses cardiomyocyte hypertrophy.

### Setd7 overexpression triggers cardiomyocyte hypertrophy

We next examined the possible link between Setd7 expression level and development of hypertrophy by exogenously overexpressing Setd7 in cultured NRCMs. Infection of cardiomyocytes with a *Setd7* lentivirus increased the Setd7 expression (**Figure S3A**). The protein levels of ANP and BNP were significantly increased in *Setd7* lentivirus cells compared with control (**Figure S3A**). Consistently, *Nppa*, *Nppb* and *Myh7* (myosin heavy chain-β) mRNA levels (**Figure S3B**) and the cell size markedly increased as a result of *Setd7* overexpression (**Figure S3C**). Furthermore, Setd7 overexpression further enhanced hypoxia-induced cardiomyocyte hypertrophy, where ANP and BNP expression and the cell-surface area were further increased compared with the parameters in hypoxia exposed cells (**Figure 3H-J**). These in vitro results indicate that overexpressing Setd7 is also sufficient to trigger cardiomyocyte hypertrophy.

### Lipid peroxidation participates in Setd7-mediated cardiac hypertrophy

Novel researches show that ferroptosis plays an important role in the pathophysiology of cardiac hypertrophy(Wang *et al*, 2020; Yin *et al*, 2020). Lipid peroxidation is a hallmark of ferroptosis and has been implicated in various heart diseases(Park *et al*, 2019; Wang *et al*, 2022). GPx4 and the cystine/glutamate antiporter system xCT (also known as solute carrier family 7 member 11; SLC7A11) are the main functional proteins that mitigate intracellular lipid peroxidation. Their inactivation impairs intracellular glutathione (GSH) synthesis, resulting in the accumulation of lipid peroxidation products and triggering ferroptosis(Yang *et al*, 2014). GPx4 is a phospholipid hydroperoxidase that protects cells against membrane lipid peroxidation. In line with previous reports(Liu *et al*, 2023), hypoxia significantly decreased xCT and GPx4 expression compared with control. Notably, we observed that Setd7 knockdown or the addition of inhibitor significantly restored the protein level of xCT and GPx4 (**Figure 4A and B**). Subsequently, the iron ion level, malondialdehyde (MDA), a marker of oxidative stress-induced lipid peroxidation, and glutathione (GSH), an antioxidant as a cofactor for GPx4 to exert antioxidation role, were detected. Results showed that inhibition or silencing Setd7 markedly reversed the iron ion level, the content MDA and GSH (**Figure 4C**). Prostaglandin endoperoxide synthase 2 (*Ptgs2*), is commonly regarded as one of the markers of ferroptosis(Yang *et al*., 2014). In cardiomyocytes, hypoxic exposure led to a significant increase of *Ptgs2* mRNA, which was inhibited by the application of siRNA and specific inhibitor targeting Setd7 **(Figure 4D)**. Conversely, Setd7 overexpression dramatically inhibited xCT and GPx4 expressions, while induced *Ptgs2* mRNA expression (**Figure S3D and E**). Consistently, compared with hypoxia, Setd7 overexpression dramatically exaggerated hypoxia-mediated xCT and GPx4 downregulation, the iron ion level, and the MDA content (**Figure 4E and F**). Again, lipid peroxidation marked by lower level of 4-hydroxynonenal (HNE)(Park *et al*, 2021) was found in heart sections of Setd7^+/−^ mice compared with WT mice. (**Figure 4G**). Our data overall suggest that lipid peroxidation participates in Setd7-mediated cardiac hypertrophy.

**Figure 4.**
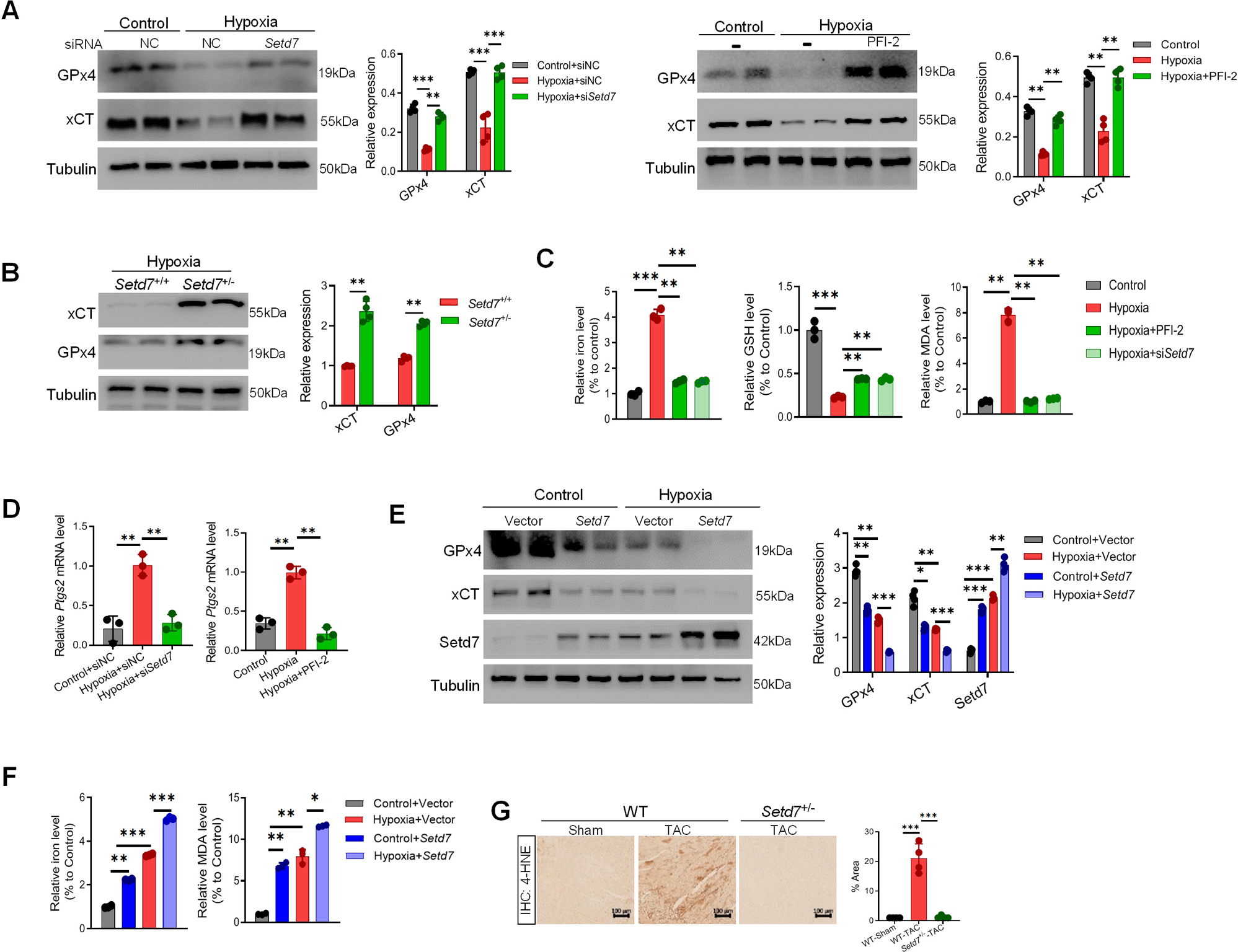
Lipid peroxidation participates in Setd7-mediated cardiac hypertrophy. (**A**) Western blot of GPx4 and xCT in NRCMs knockdown or inhibiting Setd7 upon hypoxia and quantitative analysis is on the right. (**B**) Western blot of GPx4 and xCT in neonatal *Setd7*^+/+^ and *Setd7*^+/−^ mice cardiac myocytes after hypoxia. Quantitative analysis is on the right. (**C**) Iron ion level, MDA and GSH content in hypoxia-induced NRCMs after Setd7 knockdown or inhibition by PFI-2 were assessed. (**D**) *Ptgs2* mRNA levels in NRCMs after Setd7 knockdown or inhibition by PFI-2 were determined by RT-qPCR. (**E**) NRCMs were infected with Lentivirus-*Setd7* or vector as control upon normoxia or hypoxia condition. The expressions of xCT and GPx4 were analyzed by Western blot. (**F**) Iron ion and MDA level in NRCMs infected with Lentivirus-*Setd7* or vector as control. (**G**) Representative images of heart sections stained with anti-4-HNE in WT and *Setd7*^+/−^ mice with TAC. Quantitative analysis is on the right. n=3 or 4. * represents *p* < 0.05, ** represents *p* value < 0.01, *** represents *p* value < 0.001 (Student’s t-test or One-way ANOVA).

### GPx4-mediated lipid peroxidation is responsible for Setd7-triggred cardiac hypertrophy

Our aforementioned data showed that blockade of Setd7 significantly impaired the increase in lipid peroxidation products following hypoxia exposure. Because GPx4 is known to be a main endogenous antioxidant peroxidase, we asked whether GPx4 mediated lipid peroxidation is responsible for Setd7-mediated cardiac hypertrophy. To test this possibility, Ras-selective lethal small molecule 3 (RSL3), the inhibitor of GPx4(Cheng *et al*, 2024) (**Figure S4A**), was added to Setd7 knockdown NRCMs following hypoxia exposure. We found that Setd7 knockdown preventing hypertrophy was partly abolished by RLS3, as demonstrated by increased ANP, BNP protein levels (**Figure S4B**) and cell size (**Figure S4C**). Primary cultured *Setd7*^+/+^ and *Setd7*^+/−^ cardiomyocytes were incubated with RLS3 to decrease the levels of GPx4. Subsequently, they were subjected to hypoxia and cardiac hypertrophy was then evaluated. *Setd7*^+/−^ cardiomyocytes resisted cardiac hypertrophy compared to *Setd7*^+/+^, by contrary, RLS3 partly neutralizes antihypertrophic effect of *Setd7*^+/−^ (**Figure S4D and E**). Furthermore, treatment with the ferroptosis inducer Erastin revealed that mitigated cardiomyocytes hypertrophy by Setd7 knockout partially was blocked by Erastin (**Figure S4F and G**).

To test the effects of neutralizing cellular lipid peroxides on Setd7-mediated myocytes hypertrophy, we treated overexpressing Setd7 NRCMs with lipid radical scavenger ferrostatin-1 (Fer-1). As expected, Fer-1 rectified aberrant lipid peroxidation in overexpressing Setd7 NRCMs (**Figure S5A and B**). Having established the rescue effect of Fer-1 on Setd7 overexpression-induced lipid peroxidation, we next sought to assess whether Fer-1 suppressed myocytes hypertrophy phenotypes. Notably, overexpressing Setd7 failed to induce myocytes hypertrophy and inflammatory response in the presence of Fer-1 (**Figure S5C-E**). According to the above results, Setd7 modulates cardiac hypertrophy, at least in part, through direct regulation of lipid peroxidation-dependent pathways.

### WWP2 involves in Setd7-mediated GPx4 degradation

Setd7-mediated GPx4 decrease results in the accumulation of lipid peroxidation, we sought to understand the mechanism by which Setd7 suppresses GPx4 expression. First, we inhibited protein translation by incubating the cells with cycloheximide (CHX) and determined the GPx4 protein level and stability by Western blotting. Hypoxia decreased expression of GPx4, whereas inhibiting Setd7 could maintain GPx4 abundance in the presence of CHX (**Figure 5A**). In addition, NRCMs were challenged with a proteasome inhibitor MG132, autophagic inhibitors including 3-MA and chloroquine (CQ), followed to hypoxia exposure, respectively. We found MG132 recovered the expression of GPx4 in hypoxia exposed cells **(Figure 5B)**. Consistently, hypoxia exposure enhanced the ubiquitination of GPx4, whereas the ubiquitination levels of GPx4 was blocked by Setd7 inhibition (**Figure 5C**). Altogether, these findings indicate that Setd7 promotes GPx4 degradation by the proteasome pathway. WW domain containing E3 ubiquitin protein ligase 2 (WWP2) is highly expressed in the heart. We found that hypoxia myocytes and heart exhibited up-regulation of WWP2, however, blocking Setd7 inhibited increased WWP2 expression caused by hypoxia exposure (**Figure 5D-F**). Notably, Setd7 overexpression also enhanced protein and mRNA levels of WWP2 (**Figure 5G and H**). These results demonstrate that Setd7 modulates WWP2 expression.

**Figure 5.**
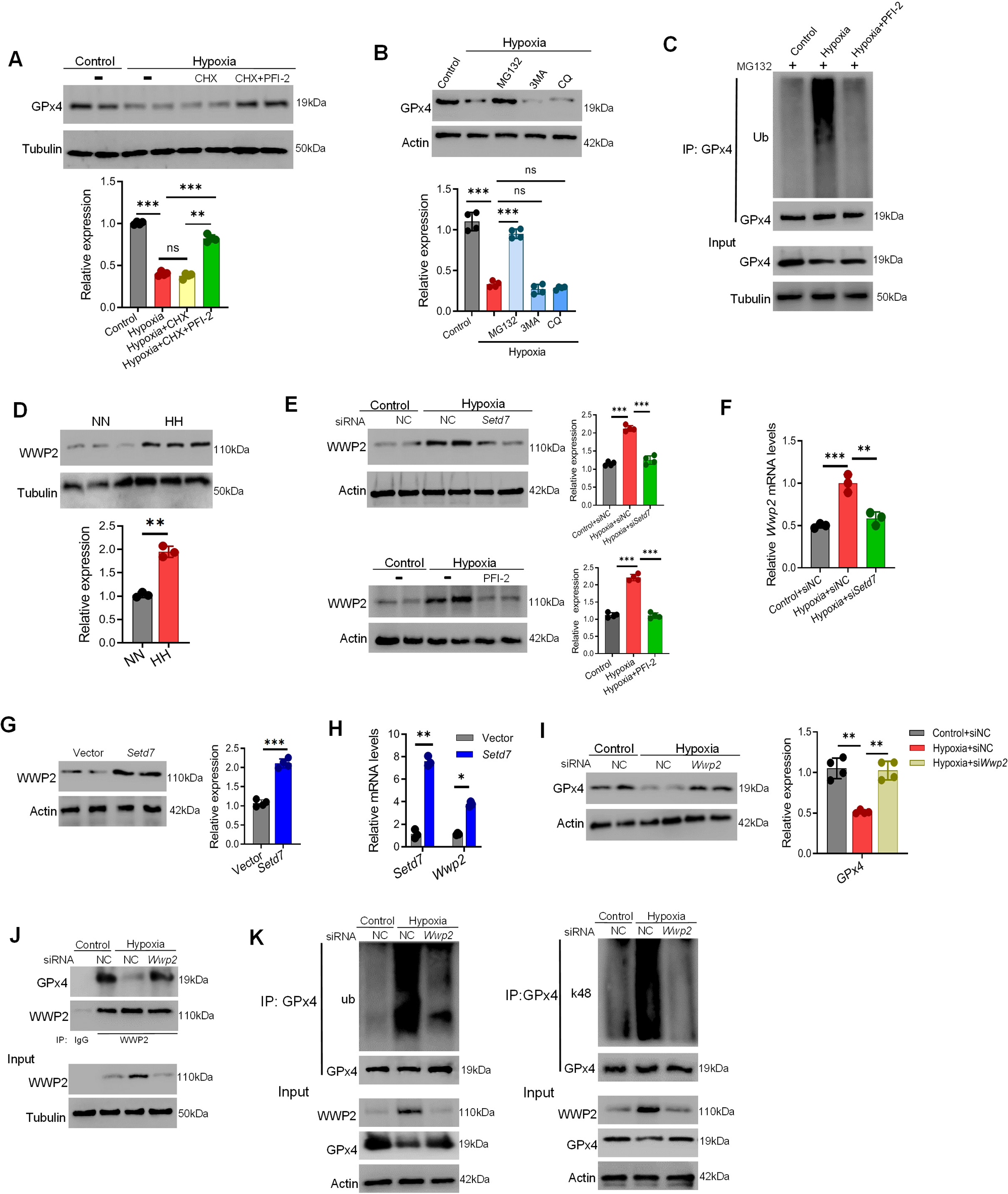
WWP2 involves in Setd7-mediated GPx4 degradation in hypoxic NRCMs. (**A**) The effect of Setd7 inhibitor PFI-2 on the expression of GPx4 in response to hypoxia followed by treatment with 100 μM CHX for 12 h in NRCMs. (**B**) Western blot analysis of protein levels of GPx4 after treatment with MG132, 3MA, CQ under hypoxia. (**C**) MG132 was added prior to the collection of protein lysate. Lysates were immunoprecipitated using GPx4 antibody and subsequently immunoblotted with antibodies against ubiquitination. (**D**) Expression of WWP2 was assessed in heart tissue of normoxia (NN) and hypobaric hypoxia (HH) mice. (**E**) The protein level of WWP2 was assessed in NRCMs under normoxia or hypoxia with or without Setd7 knockdown or inhibition. (**F**) mRNA levels of WWP2 were assessed in NRCMs under normoxia or hypoxia with or without Setd7 knockdown. (**G**) Expression of WWP2 was assessed in Setd7 overexpression NRCMs. (**H**) The Wwp2 mRNA increased in Setd7 overexpression NRCMs. (**I**) GPx4 protein expression before and after knockdown of WWP2 compared to control siRNA (NC) in NRCMs upon hypoxia. (**J**) Validation of the interaction between WWP2 and GPx4 in NRCMs by Co-IP. (**K**) Lysates were immunoprecipitated using anti-GPx4 and subsequently immunoblotted with antibodies against ubiquitination or k48 in NRCMs infected with si*Wwp2* or siNC. n=3 or 4. * represents *p* < 0.05, ** represents *p* < 0.01, *** represents *p* < 0.001, ns represents no significance (Student’s t-test or One-way ANOVA).

WWP2 belongs to the NEDD4 family of E3 ubiquitin protein ligases, we further found WWP2 knockdown recovered GPx4 protein level compared to hypoxia exposure (**Figure 5I)**. Meanwhile, co-immunoprecipitation results showed that the interaction between exogenous WWP2 and GPx4 was significantly enhanced upon WWP2 knockdown (**Figure 5J**). To further assess GPx4 ubiquitination induced by WWP2, we knocked down WWP2 along with overexpression ubiquitin in HEK293T cells. The ubiquitin-conjugated GPx4 proteins were pulled down. Western blotting showed that knockdown of WWP2 obviously decreased GPx4 ubiquitination (**Figure 5K**). These results demonstrate that WWP2 interacts with GPx4 and promotes GPx4 polyubiquitination degradation via the ubiquitin-proteasome pathway, thus WWP2 is dispensable to GPx4 protein stability.

Next, we confirmed the role of WWP2 in hypoxia-induced cardiac hypertrophy, and found WWP2 knockdown inhibited hypoxia-induced cardiac hypertrophy by decreasing the cell surface area (**Figure S6A**) and by downregulating the levels of ANP and BNP (**Figure S6B and C**). These results confirm that WWP2-mediated GPx4 degradation was involved in hypoxia-induced cardiac hypertrophy.

### Setd7 modulates WWP2 expression by E2F1 transcription factor

The above results suggest that Setd7-mediated cardiac hypertrophy via WWP2 promoting GPx4 degradation. Next, the detailed mechanism of WWP2 up-regulation by Setd7 was investigated. E2F1 (E2 promoter binding factor 1) is a transcription factor, which has strong correlation with Setd7 predicted by HitPredict database (**Figure 6A**), playing important roles in cardiac remodeling and metabolic reprogramming of cardiomyocytes(Chou *et al*, 2022; Dassanayaka *et al*, 2019). We found hypoxia induced E2F1 expression, as demonstrated by immunoblotting and fluorescence intensity (**Figure 6B and C**), and Setd7 depletion significantly decreased E2F1 expression (**Figure 6D and E**). Consistently, Setd7 overexpression also enhanced E2F1 expression (**Figure 6F**). Because Setd7 methylates histone 3 lysine 4 (H3K4) in the enhancer or promoter regions associated with transcriptional activation, we also examined H3K4me2/3 expression. Consistent with Setd7 level, hypoxia induced H3K4me2/3 expression, but Setd7 depletion significantly decreased H3K4me2/3 level (**Figure S7**). Further, ChIP assays were performed in NRCMs exposed hypoxia. We found that Setd7 and Setd7-catalyzed H3K4me2/3 were occupied on the promoter of E2F1, however, blocking Setd7 reduced Setd7 and H3K4me1/2 abundance on the promoter of E2F1 (**Figure 6G**). Subsequent validation confirmed that Setd7 indeed influenced alterations in *E2f1* transcriptional levels (**Figure 6H**). These results suggest that Setd7-catalyzed H3K4me2/3 are involved in hypoxia induced E2F1 expression.

**Figure 6.**
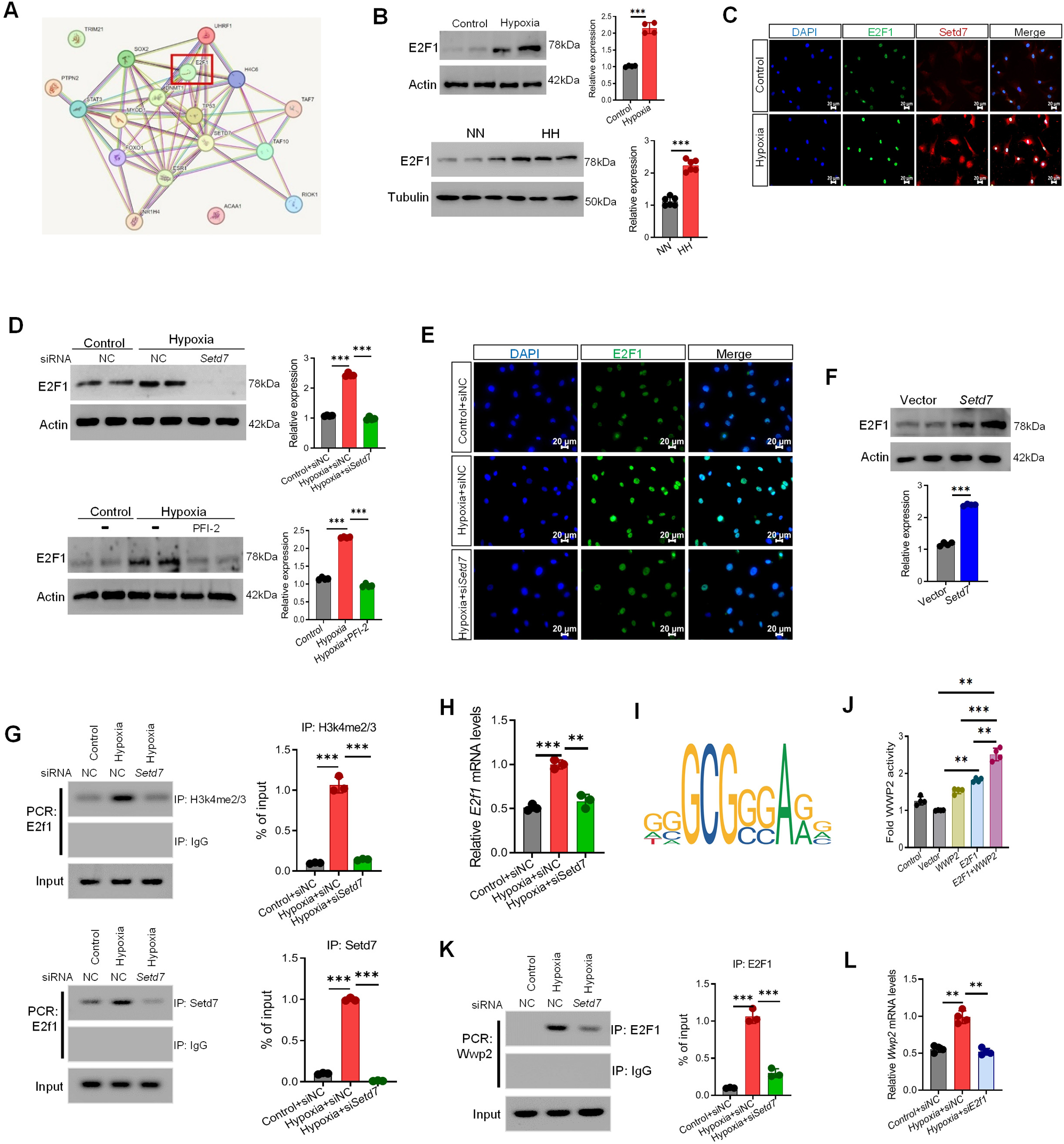
Setd7 modulates WWP2 expression by transcription factor E2F1. (**A**) The network of top 20 proteins with a strong correlation with Setd7 was constructed by String based on a gene set derived from HitPredict. (**B**) Western blotting analysis of E2F1 expression in NRCMs and heart tissue under normoxia or hypoxia condition. (**C**) Immunofluorescence double staining of E2F1 and Setd7 in NRCMs treated with or without hypoxia. Scale bar: 20 μm. (**D**) The protein level of E2F1 in NRCMs in knocking down or inhibiting *Setd7* NRCMs upon hypoxia. (**E**) Immunofluorescence staining of E2F1 in NRCMs infected with siRNA *Setd7* or control (siNC) upon normoxia or hypoxia condition. (**F**) The protein levels of E2F1 in NRCMs upon *Setd7* overexpression. (**G**) The ChIP-PCR results showed the abundance of H3K4me2/3 and Setd7 on the promoter of E2F1 in hypoxia NRCMs transfected with siNC or si*Setd7*. Quantitative analysis is on the right. (**H**) The mRNA levels of *E2f1* when knocking down the Setd7 in NRCMs under normoxia or hypoxia condition. (**I**) Motif of E2F1 that can bind to the *Wwp2* promoter region predicted by JASPAR. (**J**) Luciferase reporter assay demonstrating the effect of transcription factor E2F1 on the promoter activity of *Wwp2*. Luciferase activity was measured and normalized to the Renilla luciferase control. (**K**) ChIP analysis of E2F1 binding to the promoter of *Wwp2* in hypoxia NRCMs transfected with siNC or siSetd7. (**L**) The mRNA levels of *Wwp2* when knocking down the E2F1 in NRCMs under normoxia or hypoxia condition. n=3, 4 or 6. ** represents *p* < 0.01, *** represents *p* < 0.001 (Student’s t test or one-way ANOVA).

Furthermore, we predicted the transcription factors that can bind to the WWP2 promoter region using the PROMO database, and found that E2F1 does have a binding site on the WWP2 promoter and then verified its binding sequence using the JASPAR database (**Figure 6I**). To examine WWP2 transcriptional activity in vitro, we constructed a luciferase reporter gene containing the DNA-binding consensus sequence. We then performed reporter assays where E2F1 was co-transfected in HEK293T cells along with the reporter gene WWP2. We observed that WWP2 alone had faint transcriptional activity, but E2F1 co-expression significantly augmented it (**Figure 6J**). Meanwhile, E2F1 occupied at the promoter of *Wwp2* in myocytes exposed to hypoxia, blocking Setd7 reduced E2F1 abundance at the promoter (**Figure 6K**), and E2F1 knockdown also inhibited *Wwp2* mRNA levels (**Figure 6L**).

In addition, to further confirm the role of E2F1 in hypertrophy, we transfected NRCMs with siRNA against E2F1. Firstly, silencing E2F1 indeed inhibited the expression of WWP2 (**Figure S8A**), meanwhile, hypoxia-induced cardiomyocytes hypertrophy was abrogated by E2F1 knockdown, as indicated by the increase in hypertrophic marker gene expression (ANP, BNP) (**Figure S8A and B**) and cell size (**Figure S8C**). Further validation revealed that Setd7 overexpression induced WWP2 and 4-HNE expression, cardiomyocyte hypertrophy phenotype, and these effects were abrogated by E2F1 knockdown (**Figure S8D-F**). The results reveal that Setd7 modulates WWP2 expression by E2F1 transcription factor.

## Discussion

Pathological cardiac hypertrophy is closely related to increased risk of chronic heart failure. Setd7 is a member of the protein lysine methyltransferases family, which is reported to regulating various pathological and physiological reaction. This study is the first, to our knowledge, to characterize a crucial role for Setd7 in pathological cardiac hypertrophy and remodeling. We observed that Setd7 is significantly elevated in pathological hypertrophy stimuli cardiomyocytes and mouse failing hearts. Subsequently, we found that mice lacking Setd7 remarkably preserved cardiac function after TAC, whereas Setd7 overexpression in cardiomyocytes deteriorated hypertrophy phenotype. Further in vitro analyses revealed that Setd7 mediated E2F1 activation induces WWP2 expression to promote GPx4 ubiquitination and degradation, resulting in widespread lipid peroxidation and boosting pathological cardiac hypertrophy. Collectively, our data identify a novel and important role of Setd7 in pathological cardiac hypertrophy and suggest that targeting Setd7 is a promising therapeutic strategy to attenuate pathological cardiac hypertrophy and heart failure.

Setd7 is a member of the protein lysine methyltransferases family. It contains a SET domain. Recent studies demonstrate that Setd7 regulates cellular pathways involved in various diseases through specifically methylating both lysine H3K4 as well as a number of non-histone substrates, including p53, E2F1, RelA, and other important transcription factors (Ea & Baltimore, 2009; Kontaki & Talianidis, 2010; Kouskouti *et al*, 2004; Wang *et al*, 2001) (Chuikov *et al*, 2004). However, whether and how hypertrophic stress could regulate expression of Setd7 in cardiomyocytes was unclear. In the present study, we first observed that the expression level of Setd7 was markedly increased in cardiomyocytes hypertrophy in response to pathological hypertrophy stimuli (hypoxia and phenylephrine) and pressure overload. We then examined the development of pathological cardiac hypertrophy using genetic Setd7 ablation mice. In the experimental TAC model, mice lacking Setd7 manifested markedly reduced pathological myocardial hypertrophy and developed less interstitial fibrotic changes during chronic stress loading. Meanwhile, Setd7 ablation significantly preserved cardiac function and reduced ANP, BNP and MHC expression in hearts after. Consistently, knockdown or elective Setd7 inhibitor also exerted antihypertrophic effect in hypoxia-induced hypertrophy in vitro model, highlighting that reducing Setd7 has an antihypertrophic effect against pathological cardiac hypertrophy.

Higher levels of oxidative stress with subsequent accumulation of ROS and lipid peroxidation have been commonly implicated in pathological cardiac hypertrophy(Alzahrani *et al*, 2021; Baba *et al*, 2018). Consistent with previous reports, we showed that levels of MDA and 4-HNE, products of lipid peroxidation were significantly increased in hearts after TAC injury or exposed to hypoxia with normal heart. Hypoxia exposure inhibited GPx4 expression and induced lipid peroxidation, inhibiting GPx4 clearly enhanced lipid peroxidation with concomitantly increased cardiac hypertrophy. Interestingly, we observed that Setd7 knockdown significantly recovered GPx4 expression and inhibited lipid peroxidation in hypoxia cardiomyocyte. Notably, in the conditions of ferroptosis inducer RSL3 or Erastin supplementation, Setd7 knockdown failed to exert antihypertrophic effect, suggesting that GPx4 expression in cardiomyocyte is necessary to mediate the benefit of Setd7 knockdown. On contrary, lipid radical scavenger ferrostatin-1 rectified aberrant lipid peroxidation and cardiomyocyte hypertrophy in overexpressing Setd7 NRCMs. Our findings support the notion that Setd7 knockdown inhibited lipid peroxidation via recovered GPx4 may have an important role in modulating cardiac hypertrophy development, especially in the setting of reduced GPx4 expression like pathological cardiac hypertrophy.

GPx4, a critical regulator of ferroptosis, plays a pivotal role in mediating lipid peroxidation-dependent, non-apoptotic cell death. Conditional knockout studies of GPx4 in embryonic fibroblasts have demonstrated its essential function in this process(Seiler *et al*, 2008). As a negative regulatory element of ferroptosis, numerous bioactive molecules have been identified that target GPx4 to induce ferroptosis(Sun *et al*, 2023b). The activity and stability of GPx4 are modulated by various factors. Persistent oxidative stress and concomitant GSH deficiency lead to the formation of dehydroalanine during β-cleavage, resulting in the irreversible inactivation of GPx4(Orian *et al*, 2015). Additionally, post-translational modifications of GPx4, particularly ubiquitination, have received increasing scientific attention due to their critical role in regulating GPx4 stability. Numerous ubiquitination enzymes, including STUB1(Sun *et al*., 2023b), TRIM26(Wang *et al*, 2023), and TRIM21(Sun *et al*, 2023a), have been identified as key regulators in various diseases through their modification of GPX4. These enzymes play critical roles in the ubiquitination-mediated regulation of GPx4, thereby influencing its stability and activity, and consequently impacting disease progression.

WWP2, an E3 ubiquitin ligase, participates in cellular processes by targeting various substrates for degradation. It is highly expressed in the heart and implicated in various myocardial pathological processes(Chen *et al*, 2024b). Studies have shown that WWP2 mediates the ubiquitination and degradation of PARP1, thereby regulating isoproterenol induced cardiac remodeling and exerting cardioprotective effects(Zhang *et al*, 2020). Additionally, LINC01588 interacts with HNRNPL to mitigate hypoxia/reoxygenation-induced cardiomyocyte injury by promoting WWP2-mediated SEPT4 degradation(Song *et al*, 2022). We found in the present study that enhanced WWP2 expression by Setd7 could interact with GPx4 and promote ubiquitination and degradation of GPx4, leading to lipid peroxidation in hypoxic hypertrophic cardiomyocytes. This study identified GPx4 as a new substrate of WWP2. Previous systematic genetic analyses revealed that WWP2 upregulates extracellular matrix genes, suggesting its involvement in the regulatory network of pro-fibrotic genes in heart disease and myocardial fibrosis(Chen *et al*, 2019). Consistently, myocardial fibrosis-related markers (Tgfβ1) increased during TAC-induced cardiac hypertrophy, which suggests that WWP2 may also play a pro-fibrotic role in this model. In addition, whether WWP2 can regulate other related proteins involved in hypoxia-induced cardiac hypertrophy needs further study.

E2F1 as an important transcription factor plays a crucial role in various heart diseases through its diverse regulatory functions(Ertosun *et al*, 2016). Research has demonstrated that E2F1 knockout mice can improve myocardial infarction-induced cardiac dysfunction and ventricular remodeling(Dassanayaka *et al*., 2019). Furthermore, E2F1 can upregulate VEGF and PIGF through both p53-dependent and -independent pathways, facilitating neovascularization in the myocardial infarction area(Wu *et al*, 2014). E2F1 also contributes to the differentiation of human cardiac fibroblasts and promotes Tgfβ1-induced cell differentiation, leading to myocardial fibrosis by directly targeting CCNE2(Liao *et al*, 2021). Importantly, E2F1 has strong correlation with Setd7 predicted by HitPredict database. Moreover, transcription factor binding site predictions revealed that E2F1 can occupy specific sites within the WWP2 promoter region. Hence, we speculate that E2F1 may involve in Setd7 mediated WWP2 expression during the cardiac hypertrophy. Indeed, we discovered that Setd7 and Setd7-catalyzed H3K4 methylation bind to the promoter region of E2F1 in hypertrophic cardiomyocytes, thereby activating E2F1 transcription. Subsequently, we also observed E2F1 is occupied on the promoter of Wwp2 in myocytes exposed hypoxia to promote WWP2 transcription, blocking Setd7 delated E2F1 abundance on the promoter of Wwp2, thus inhibiting Wwp2 mRNA levels. These findings highlight the intricate regulatory mechanisms involving Setd7, E2F1, and WWP2 in cardiac hypertrophy.

This study illustrated the role of Setd7 in cardiomyocytes during pathological cardiac hypertrophy and elucidates its underlying mechanisms. However, there are still some limitations that need to be addressed. First, the mechanistic investigations were conducted in vitro, performing an in vivo experiment would further be needed to solidify these findings. Additionally, systemic knockout or overexpression of specific genes usually triggers adaptive changes due to compensatory functions, which may lead to some potential misunderstandings of the results. In the study, we utilized systemic Setd7 knockdown mice. We are aware that the protective effects of Setd7 deficiency may not be exclusively contributed by other organs or cells. It is possible that the results currently observed in vivo may be attributed to the action of multiple factors, which would require further experiments to confirm.

In summary, this study reveals the Setd7 directly enhances E3 ubiquitin ligase WWP2 expression through enriching the transcriptionally activation marks H3K4 on the E2F1 promoters, resulting in promoting polyubiquitination and degradation of GPx4, and then exacerbating lipid peroxidation and the development of cardiac hypertrophy. Blocking Setd7 mediates the myocardial protective effect against pathological cardiac hypertrophy through E2F1-WWP2-GPx4 axis. Our data demonstrates a new and previous unrecognized role of Setd7 in cardiac hypertrophy development.

## Abbreviations

293T: human embryonic kidney 293 cells, contains the SV40 T-antigen
3-MA: 3-methyladenine
ANP: natriuretic peptide A
α-SMA: α-smooth muscle actin
BNP: natriuretic peptide B
BW: body weight
ChIP: chromatin immunoprecipitation
CHX: cycloheximide
Co-IP: Co-immunoprecipitation
DAPI: diaminidine phenyl indole
DMEM: Dulbecco’s modified Eagle medium
EF: ejection fraction
FBS: fetal bovine serum
FS: fractional shortening
GPx4: glutathione peroxidase 4
GSH: glutathione
H&E: hematoxylin and eosin
H3K4: histone H3 lysine 4
HF: heart failure
HW/TL: heart weight/tibia length
HW: heart weight
IHC: immunohistochemistry
KO: knockout
LOOH: Lipid hydroperoxides
LVID: LV internal dimension
MDA: malondialdehyde
MYH7: muscle β-isoform
NMCMs: neonatal mice cardiac myocytes
NRCMs: neonatal rat cardiomyocytes
PE: phenylephrine
Ptgs2: prostaglandin-endoperoxide synthase 2
PTM: post-translational modification
ROS: reactive oxygen species
RT-qPCR: Real time and quantitative PCR
RVH: right ventricular hypertrophy
SD: Sprague-Dawley
Setd7: SET domain containing 7
siRNA: small interfering RNA
SLC7A11 / xCT: solute carrier family 7 member 11 / system Xc-
TAC: transverse aortic construction
TL: tibia length
WGA: wheat germ agglutinin
WT: wild type

## Acknowledgments

The working model of this study was drawn using Biorender (www.biorender.com).

## Ethics declarations

All animals and the experimental protocol conformed to the Animal Welfare Act Guide for Use and Care of Laboratory Animals, and were approved by Institutional Animal Care and Use Committee (IACUC), Fudan University, China.

## Author contributions

Haibi Su and Jinghuan Wang performed experiments, analyzed the data and wrote the manuscript. Yuyu Zhang and Jie Xu contributed to in vitrio experiments. Jiayao Liu, Yuhui Li, Chenxi Xiao and Caiyun Wang contributed to the animal experiments. Xinhua Liu and Jun Chang designed the experiments and supervised the project. All authors read and approved the final manuscript.

## Data Availability

All data in this study are provided in the paper and Supplementary Materials and Tables.

---

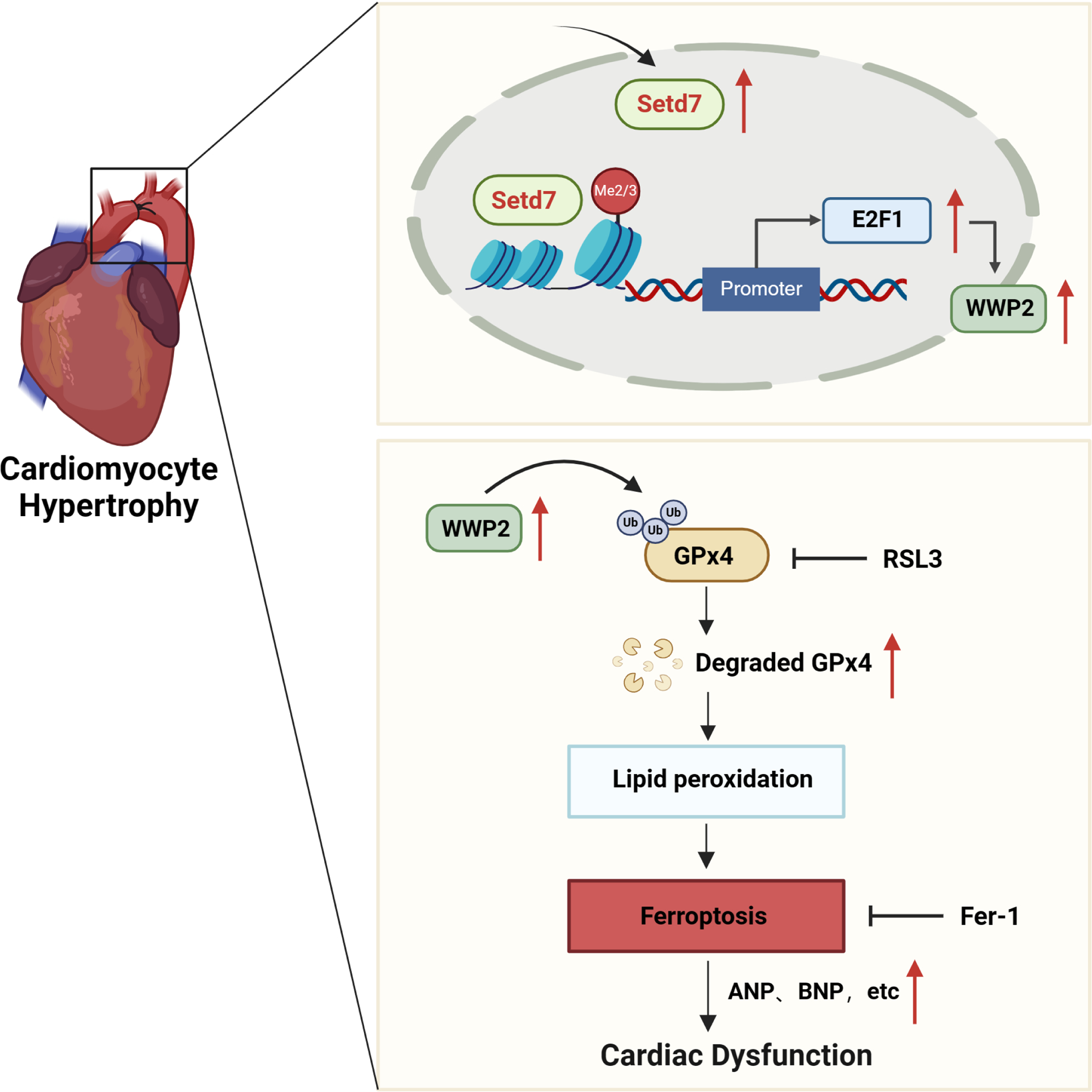

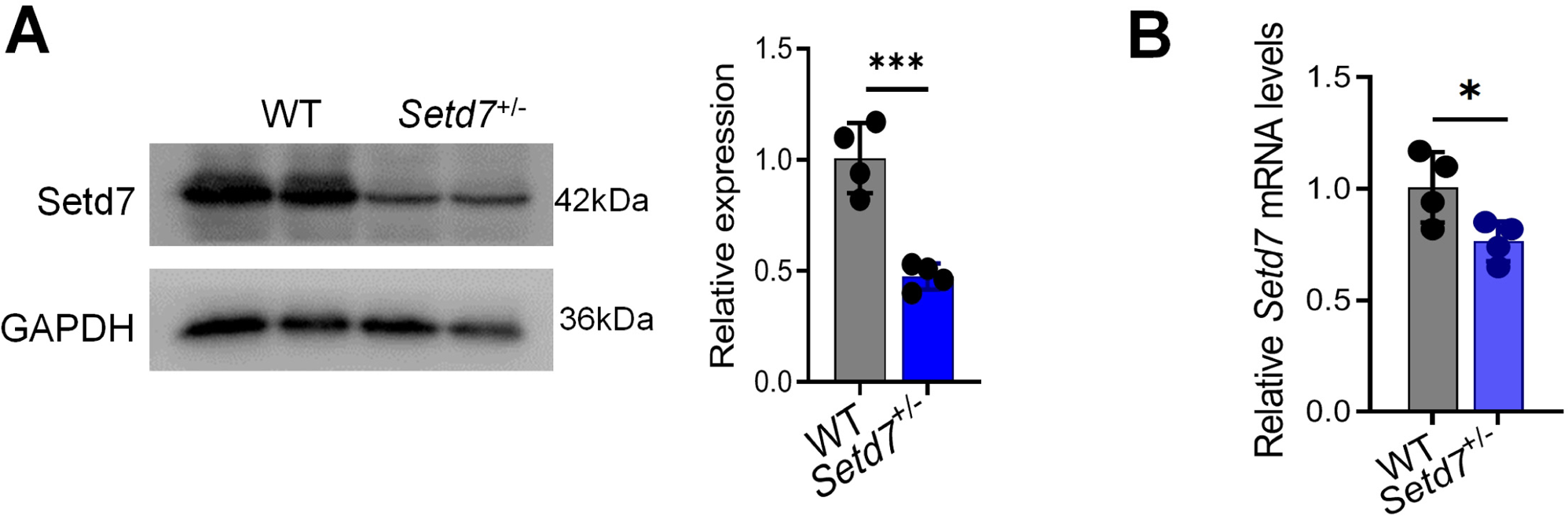

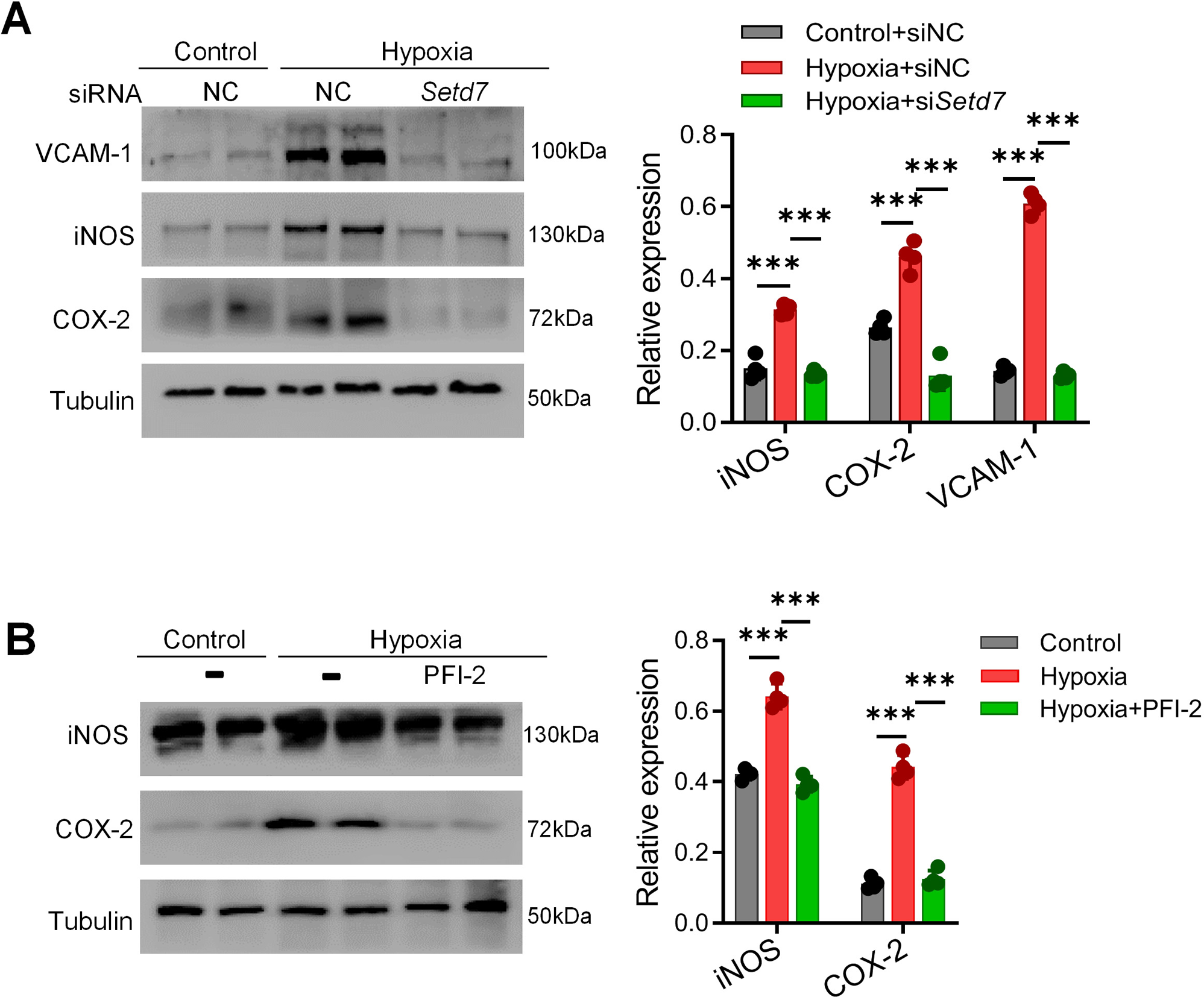

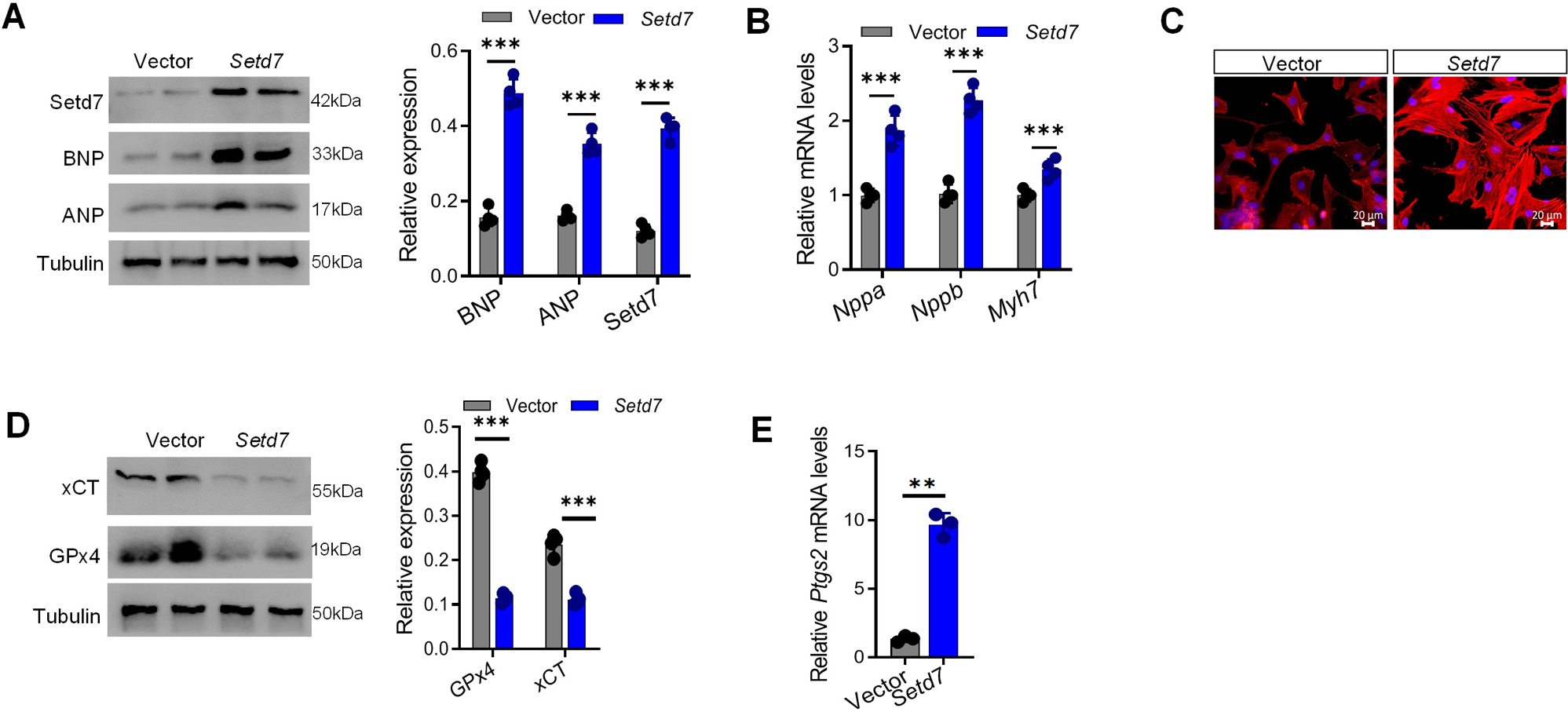

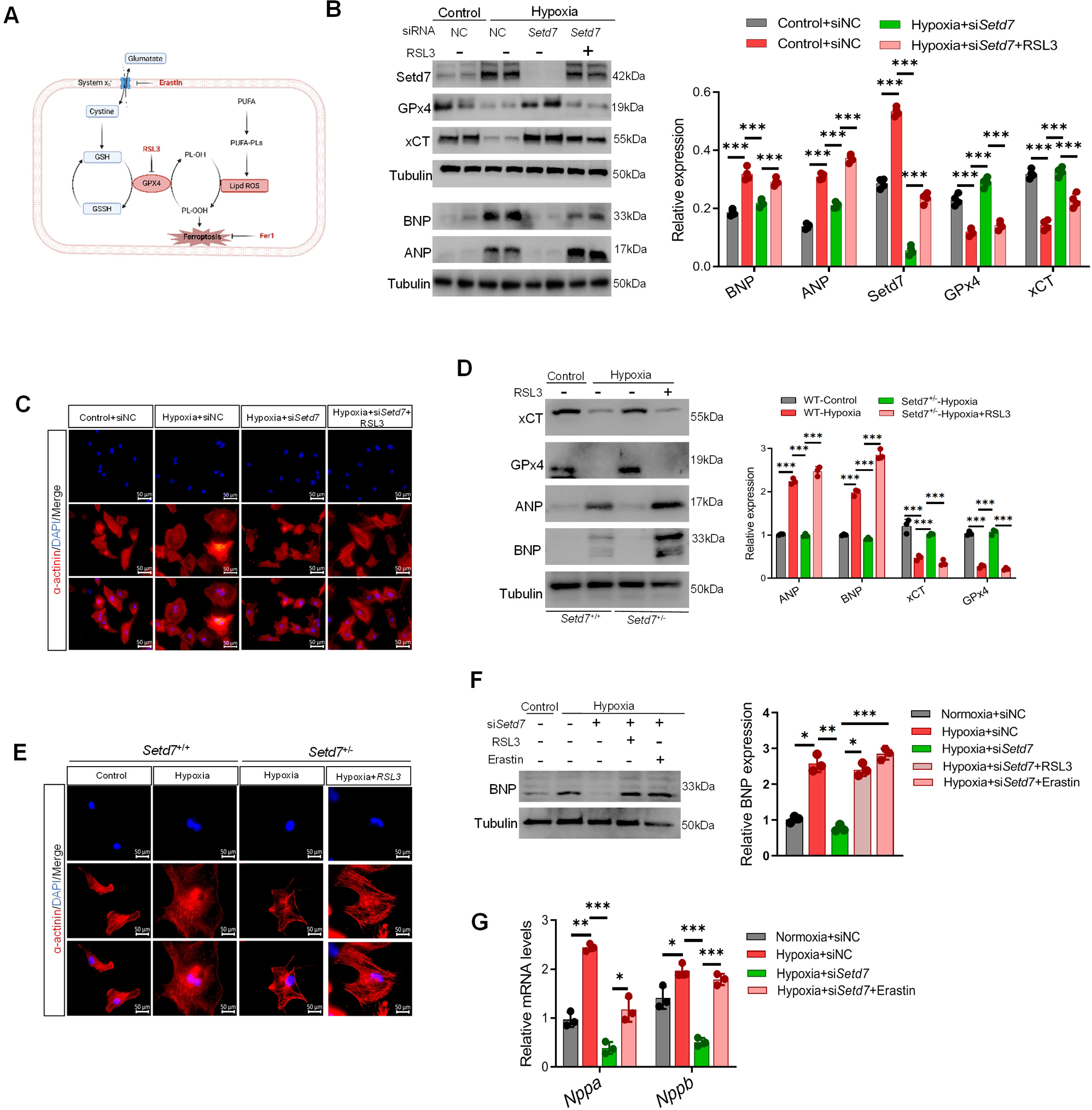

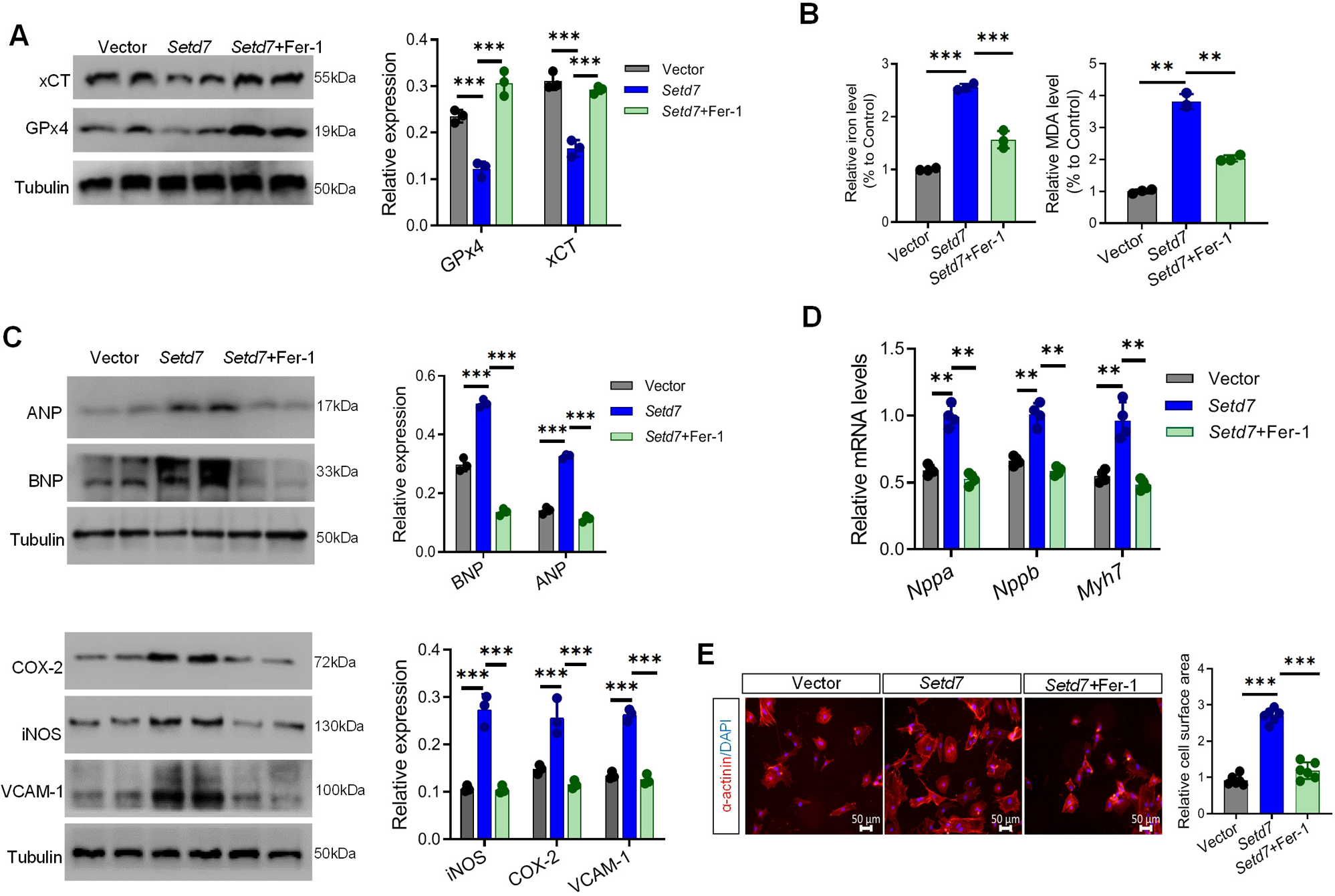

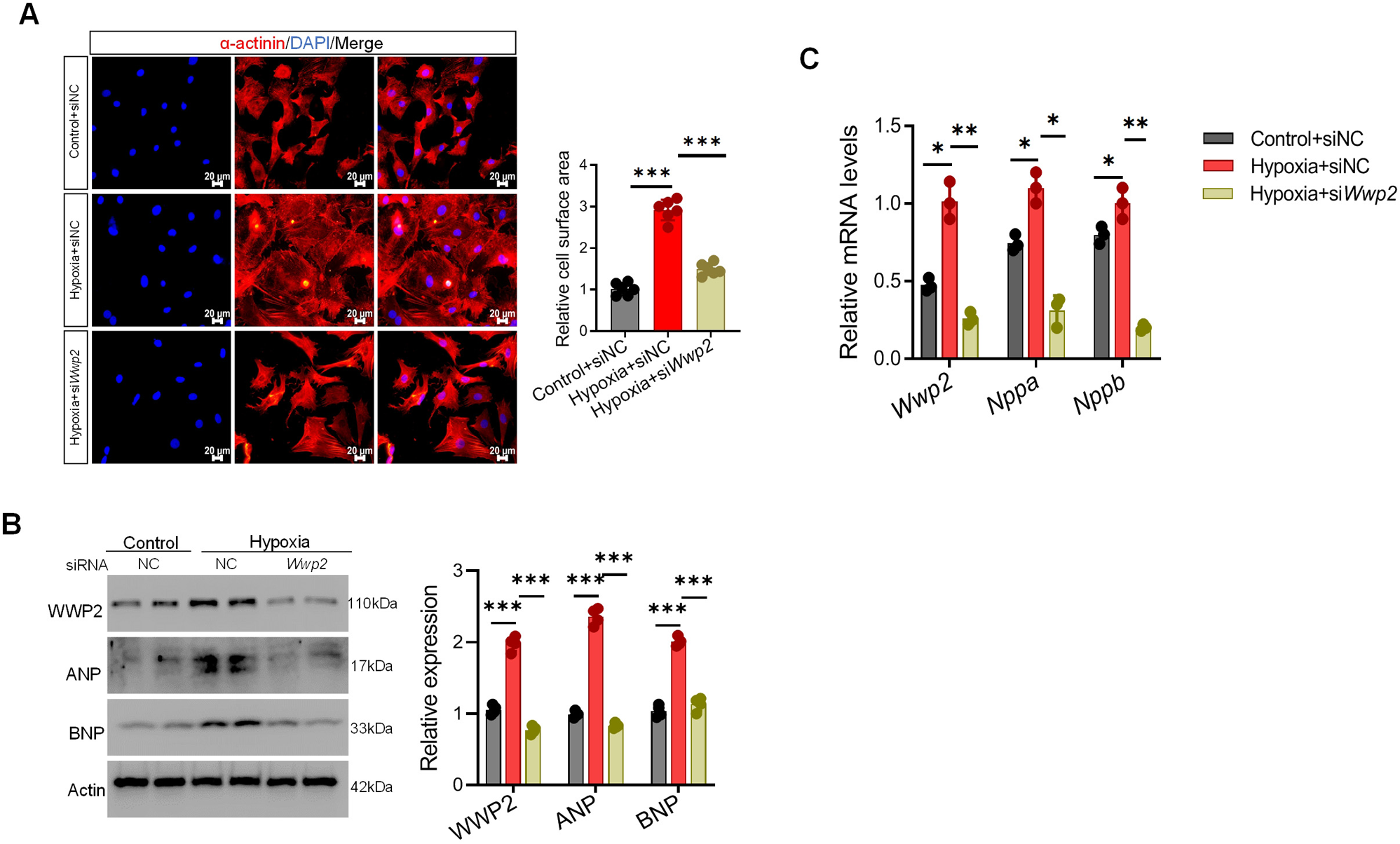

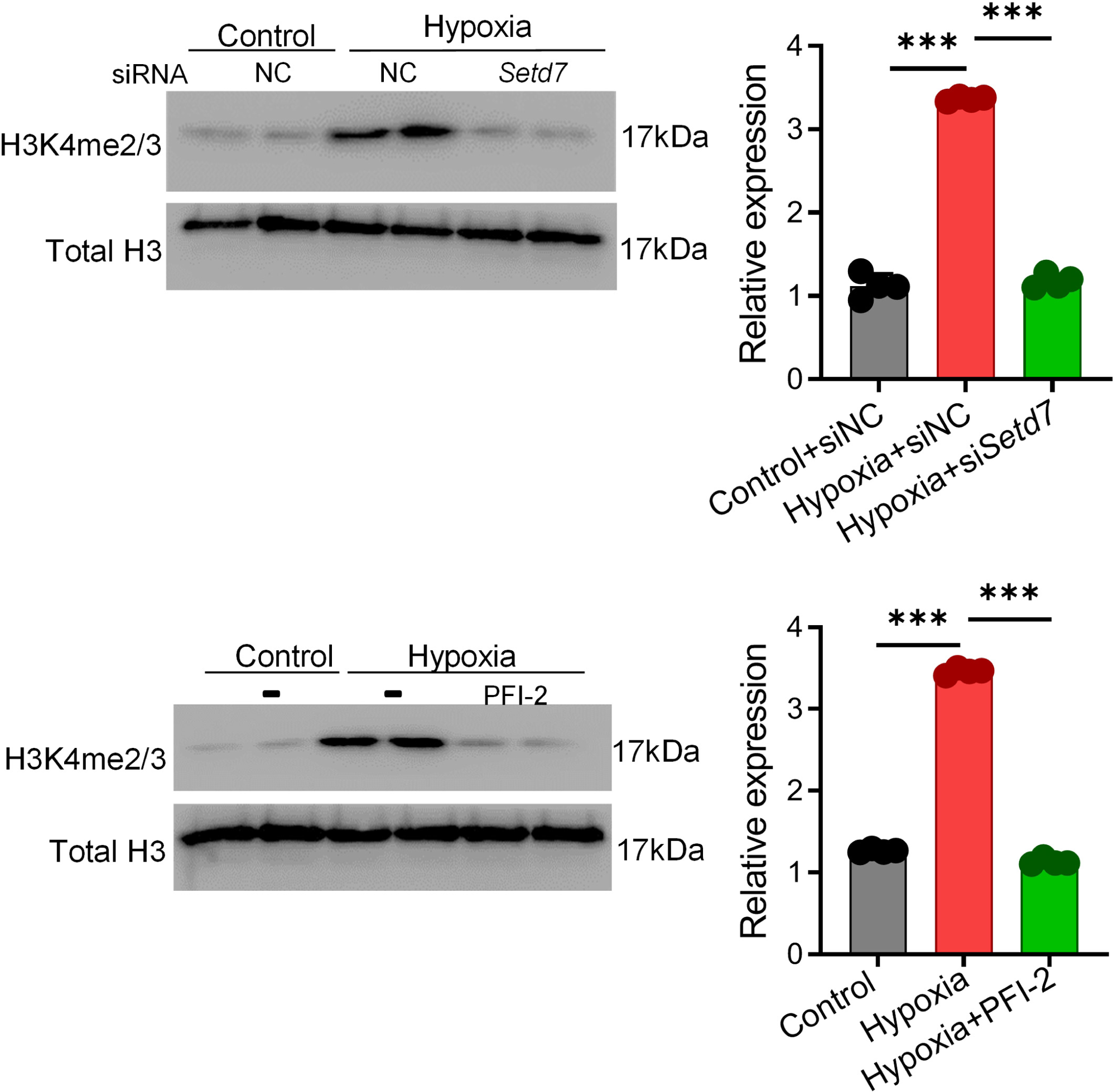

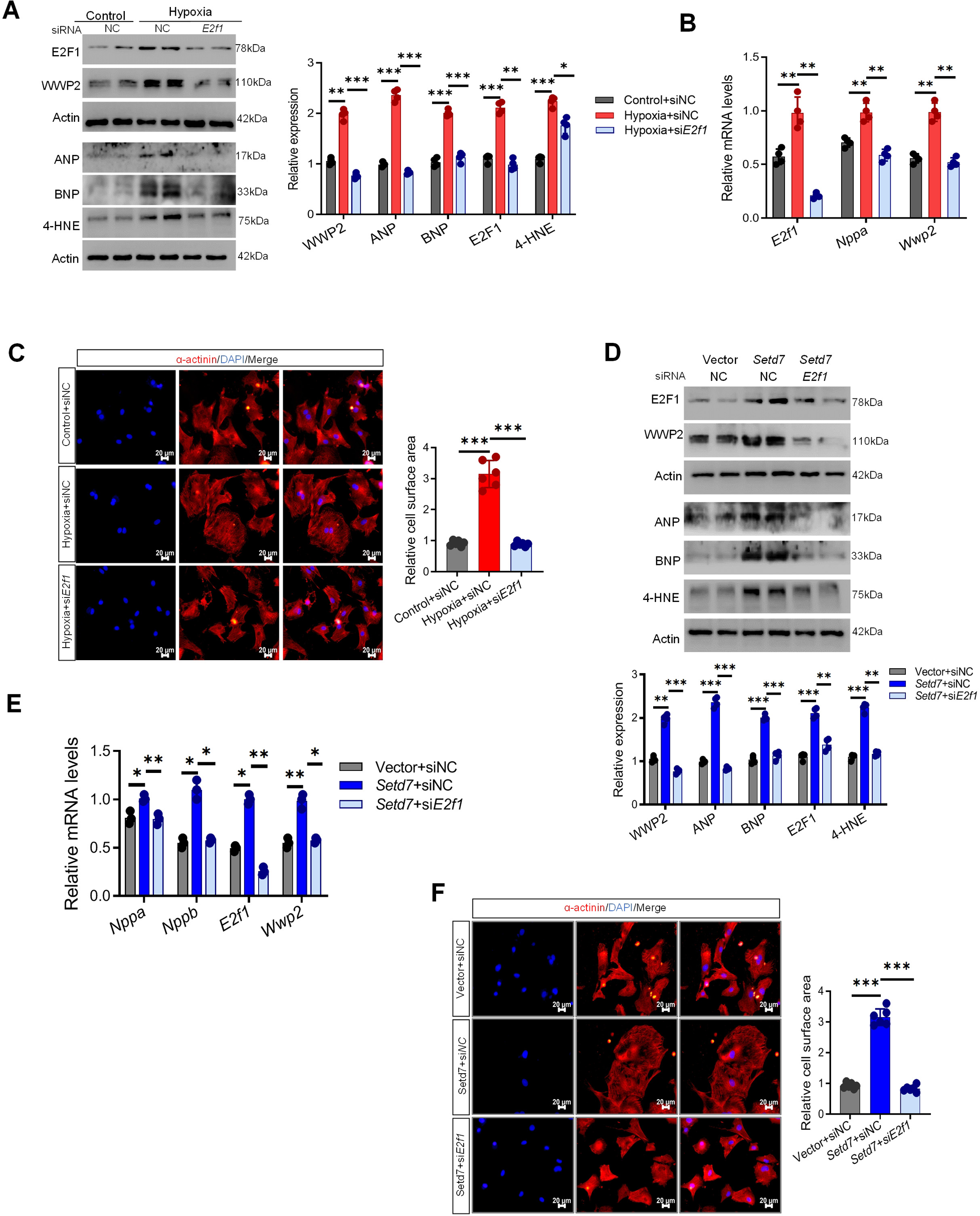

